# Biomedical Concept Recognition Using Deep Neural Sequence Models

**DOI:** 10.1101/530337

**Authors:** Negacy D. Hailu, Michael Bada, Asmelash Teka Hadgu, Lawrence E. Hunter

## Abstract

**Background:** the automated identification of mentions of ontological concepts in natural language texts is a central task in biomedical information extraction. Despite more than a decade of effort, performance in this task remains below the level necessary for many applications.

**Results:** recently, applications of deep learning in natural language processing have demonstrated striking improvements over previously state-of-the-art performance in many related natural language processing tasks. Here we demonstrate similarly striking performance improvements in recognizing biomedical ontology concepts in full text journal articles using deep learning techniques originally developed for machine translation. For example, our best performing system improves the performance of the previous state-of-the-art in recognizing terms in the Gene Ontology Biological Process hierarchy, from a previous best F1 score of 0.40 to an F1 of 0.70, nearly halving the error rate. Nearly all other ontologies show similar performance improvements.

**Conclusions:** A two-stage concept recognition system, which is a conditional random field model for span detection followed by a deep neural sequence model for normalization, improves the state-of-the-art performance for biomedical concept recognition. Treating the biomedical concept normalization task as a sequence-to-sequence mapping task similar to neural machine translation improves performance.

## Introduction

The automated identification of mentions of ontological concepts in natural language texts is a central task in biomedical information extraction. As demonstrated in BioCreAtIvE challenge evaluations^1^ and recent publications (reviewed below), despite more than a decade of effort, performance in this task remains below the level necessary for many applications. Recently, applications of deep learning in natural language processing have demonstrated striking improvements over previously state-of-the-art performance in many related natural language processing tasks (see, e.g. [42]). Here we demonstrate similarly striking performance improvements in recognizing biomedical ontology concepts in full-text journal articles using deep learning techniques developed for machine translation.

This task has often been divided into two stages: First, spans of text that refer to some ontological concept are identified. After such text spans are detected, a second stage then maps these spans to specific ontological concepts. The first step is often referred to in the literature as “named entity recognition” or “mention detection,” and the second step is often called “concept normalization.” However, as there are also previous publications that refer to either or both tasks together with similar terms, to be clear here we will call the first task “span detection,” the second task “normalization,” and the end-to-end combination of the two “concept recognition.”

We have obtained dramatic performance improvements by reconceiving concept normalization as a kind of language translation task, from ambiguous natural language into the unambiguous language of ontology terms. We used our own implementation of deep neural sequence models as depicted in Figure 2 that we developed based on coursework from https://www.deeplearning.ai/ and on the popular Open Neural Machine Translation system (OpenNMT, http://opennmt.net/). These models employ encoder-decoder architectures with attention mechanisms, a widely used approach for neural machine translation.

The Colorado Richly Annotated Full-Text (CRAFT) corpus [30] was used to train and test our models. CRAFT is a collection of 67 full-text articles largely focused on the laboratory mouse and marked up with very extensive syntactic and semantic annotations. Concept normalization tasks are defined by the set of concepts to be recognized. The most recent version of CRAFT (v3.0) contains manually curated “gold standard” annotations of text to concepts from ten different Open Biomedical Ontologies (OBOs) [40], covering a wide range of concepts seen throughout the biomedical literature, including chemical entities, biological sequence features, genes and gene products, cells, subcellular components and regions, anatomical entities, biological taxa and organisms, chemical reactions, and biological processes. Our deep learning approach achieved significant performance improvements for nine out of ten of these ontologies. The breadth of ontologies with improved performance suggests the approach should further generalize beyond the ontologies tested here.

Automated concept recognition is difficult for a variety of reasons. Human language is extremely variable, so there are many different ways to express any particular concept. For example, the Gene Ontology concept “biological regulation” (GO:0065007) is annotated to 34 different textual expressions in the latest version of CRAFT, including “modulate,” “control,” “govern,” “regulatory,” as well as many morphological variants of each of those. In addition to variability, difficulty in concept recognition arises through ambiguities. For example, the word “nucleus” could refer to an atomic nucleus (CHEBI:33252), a cell nucleus (GO:0005634), or an anatomical nucleus (UBERON:0000125) among other types. The very large number of concepts in the target ontologies is also a challenge, at least for normalization through machine learning. There are nearly a million concepts in the ten ontologies combined, and even the smaller ontologies have many thousands of concepts. Treating concept recognition as a machine learning classification task leads to an impractically large multiclass classification problem, especially given the limited amount of training data available.

The current state of the art in concept recognition in biomedical texts has been achieved through dictionary lookup [4] or with hybrids of dictionary lookup and machine learning [5, 6]. The advantage of dictionary-based systems is that a training dataset is not required. In recent years, neural network systems have been demonstrated to provide significant performance increases in many natural language processing tasks [10], but the very large number of concepts in the OBOs have until now precluded their use in the normalization task.

Our main contribution is to cast the normalization task as a kind of language translation, i.e., between a natural language text and textual IDs denoting ontological classes. Rather than treating normalization as a classification task, we treat it as a task of mapping of the sequence of characters in a text string to the sequence of characters that is the appropriate ontology identifier. This approach allowed the training of a highly effective system with the data available from CRAFT.

There are several potential advantages to this approach. The primary one is that the output to be predicted is now a relatively short string of characters, not a choice among thousands of different classes. Treating the inputs as characters, rather than as tokenized words also addresses the problem of unknown or out-of-vocabulary terms, as the model learns sub-word patterns, which covers potentially many unseen terms whose fragments (character n-grams) have been seen during training [37]. Also, as we demonstrate below, there is apparently some semantic information in identifier substrings that the system can extract. It is likely that this semantic information arises due to a curation process that produces groups of related terms with sequentially adjacent identifiers.

### Related work

Previous biomedical normalization systems are based on either dictionary lookups or hybrids of dictionary lookup systems with machine learning classifiers for postprocessing. Funk et al. [4] presented a systematic evaluation of dictionary lookup systems and offered optimal system and parameter settings for the ontologies used for concept annotation of the CRAFT corpus. Campos et al. [5] described a hybrid system employing dictionary matching and machine learning system for biomedical concept normalization. Groza et al. [9] approached the task as in information retrieval using case sensitivity and information gain. Basaldella et al. [6] described a system relying on high-recall dictionary lookups followed by a high-precision machine learning classifier. We compare our results to each of these systems.

Bahdanau, et al., [39] introduced attention mechanism to encoder-decoder architecture for neural machine translation that allows the model to automatically find relevant information from a source sentence while predicting a target word, which achieved significant performance gains over previous approaches. This is a sequence-to-sequence method that maps a sequence of characters or tokens in one language into a sequence of characters or tokens in another language. Deep neural networks have demonstrated improved performance over the prior state of the art in many natural language processing tasks [10], including email auto-response [11], morphological inflection [12, 13, 14], sentence compression [15], part-of-speech tagging and named entity recognition [16], and syntactic parsing [17].

More recently, sequence-to-sequence methods have been applied to the span detection problem. Here the problem is cast a sequence tagging or labeling, where each element of a sequence is labeled with a tag. In addition to the obvious applications such as tagging words with part-of-speech tags, this approach can be used for detecting spans, where the output sequence identifies elements at the beginning (B), inside (I), or outside (O) of a span, sometimes called BIO tagging [18]. Hung et al. (2015) [20] proposed a model based on Bidirectional Long-Short Memory (Bi-LSTM) along with a conditional random field (CRF) model for BIO tagging. Hung et al. concluded that a Bi-LSTM with CRF achieved better tagging accuracy for part-of-speech tagging, chunking and span detection than a CRF alone. Ma et al. [21] proposed an end-to-end sequence-tagging model based on bi-directional LSTM-CNN-CRF. Lample et al. [22] proposed a neural architecture for span detection; its input representation used character embedding and word embeddings to feed a Bi-LSTM+CRF architecture.

There are multiple success stories of neural sequence models applied in the biomedical domain as well. Habibi et al [23] applied the BiLSTM-CRF architecture proposed by Lample et al. for span detection on wide range of datasets in the biomedical domain. They found that their model outperforms state-of-the-art methods. The same architecture was used by Gridach [24] to recognize spans of genes and proteins. Zhao et al. [25] proposed a multiple label strategy (MLS) that can replace the CRF layer of a deep neural network for detecting spans of disease names. Korvigo et al. [26] applied a CNN-RNN network to recognize spans of chemicals and Luo et al. 2018 [28] proposed attention-based bidirectional LSTM with CRF to detect spans of chemicals. Unanue et al., 2017 [29] used bidirectional LSTM with CRF to detect spans of drug names and clinical concepts, while Lyu et al. 2017 [27] proposed bidirectional LSTM-RNN model for detecting spans of a variety of biomedical concepts. However, none of these approaches also attempted the normalization step, so they did not identify which particular concept in an ontology was detected.

## Materials and Methods

Training and evaluation of models followed the traditional two-step method for concept recognition over the OBOs that combine a sequence tagging approach to detect the spans of concepts, followed by a mapping of the contents of those spans to ontology class identifiers. Bi-LSTM and CRF methods were both evaluated for span detection, and a variety of experiments on normalization showed both the superiority of the sequence-to-sequence translation approach and elucidated the elements of the approach responsible for that performance.

### Materials

The Colorado Richly Annotated Full-Text (CRAFT) corpus [30] was used to train and test our models. The corpus is a collection of 67 full-text articles with a focus (though not exclusively) on the laboratory mouse that has been extensively marked up with both gold-standard syntactic and semantic annotations. Among the former, the annotations of segmented sentences, tokens, and part-of-speech tags were used to extract features to train a CRF model for span detection. The semantic annotations have been created with classes relying on ten Open Biomedical Ontologies:

1. Chemical Entities of Biological Interest (ChEBI): compositionally defined chemical entities (atoms, chemical substances, molecular entities, and chemical groups), subatomic particles, and role-defined chemical entities (i.e., defined in terms of their use by humans, or by their biological and/or chemical behavior)
2. Cell Ontology (CL): cells (excluding types of cell line cells)
3. Gene Ontology Biological Process (GO_BP): biological processes, including genetic, biochemical/molecular-biological, cellular and subcellular, organ-and organ-system-level, organismal, and multiorganismal processes
4. Gene Ontology Cellular Component (GO_CC): cellular and extracellular components and regions; species-nonspecific macromolecular complexes
5. Gene Ontology Molecular Function (GO_MF): molecular functionalities possessed by genes or gene products, as well as the molecular bearers of these functionalities
6. Molecular Process Ontology (MOP): chemical reactions and other molecular processes
7. NCBI Taxonomy (NCBITaxon): biological taxa and their corresponding organisms; taxon levels
8. Protein Ontology (PR): proteins, which are also used to annotate corresponding genes and transcripts
9. Sequence Ontology (SO): biomacromolecular entities, sequence features, and their associated attributes and processes
10. Uberon (UBERON): anatomical entities; multicellular organisms defined in terms of developmental and sexual characteristics.

CRAFT has had three major releases. The most recent release, version 3.0, is a significant improvement from version 2.0 with regard to the concept annotations: The annotations for all eight of the ontologies used for version 2.0 of the corpus have been updated using the classes of newer versions of these ontologies, resulting in a substantial increase in the number of annotations. Additionally, extension classes of the ontologies have been created and extensively used for annotation. (A detailed discussion of the extension classes can be found at https://github.com/UCDenver-ccp/CRAFT/tree/master/ontology-concepts/README.md.) The concept annotations for each ontology are packaged into sets created without any extension classes and sets augmented with extension class annotations. Also included along with the concept annotations are original OBO ontology files and those augmented with extension classes, as well as various class mapping files. Additionally, compared to version 2.0, concept annotations using the Molecular Process Ontology (MOP) and the Uberon anatomical ontology have been created for the articles of the corpus. The concept annotation span guidelines have also been slightly modified (also discussed in the aforementioned README file). The concept annotations of version 3.0 of the corpus are now available in Knowtator 1, Knowtator 2, brat, and UIMA XMI formats (though no longer in AO RDF or Genia XML formats).

Version 1.0 was used for evaluations in previous publications (e.g., [4], [5], and [6]), so performance of the sequence-to-sequence method was also evaluated using that first release. The sequence-to-sequence approach was evaluated on version 3.0 as well. While not strictly comparable to previously published approaches, the results on version 3.0 are most indicative of performance of the method in realistic current applications. Within each of the ontology annotation sets, versions of the corpus prior to 3.0 contain nested concept annotations (i.e., annotations with spans entirely within the spans of other annotations). For example, for the following article title, there is a nesting annotation for “Pachytene Checkpoint” and a nested annotation for “Pachytene” in the Gene Ontology Biological Process annotation set:

PMID-17696610: “Mouse <u>Pachytene Checkpoint</u> 2 (Trip13) Is Required for Completing Meiotic Recombination but Not Synapsis.”

For a given pair of nesting and nested concept annotations, the nested annotation was ignored in this work; thus, for this sentence, the annotation for “Pachytene” is ignored.

Additionally, all versions of the corpus contain discontinuous annotations, i.e., annotations of two or more discontinuous text spans. For such cases, the whole continuous span of text from the beginning of the fist span to the end of the last span, including the unannotated intervening text span(s), is used for training. For example, in the sentence PMID – 17696610 “…more closely to plants than it does to the evolutionarily more closely related worms and flies.” The span “evolutionarily… related” is annotated as (SO:0000857, homologous). For training purposes the span for the ontology term was defined as “evolutionarily more closely related.”

### Methods

*Span detection* is approached as a sequence-tagging task, using the BIO (Beginning, Inside and Outside) format to label tokens in a sentence. Two approaches to this task were evaluated.

The first approach used NERsuite [32], which is a publicly available, off-the-shelf CRF implementation that has been used in multiple biomedical text mining tools, e.g., [33, 43]. Successful training of a CRF typically requires the selection of a set of input features, and for this we used words, part-of-speech tags, and shallow parses. The features within windows of three tokens before and after a given target word were used. The BIO format is used to capture syntactic information of the shallow parses (or chunks) such as noun phrase, verb phrase etc. The details of the feature extraction steps are described in Okazaki [34].

The second approach used deep learning methods based on Reimers et al [31]. This approach uses word and character embeddings as input and does not require any feature engineering steps.

Two span detection models were trained for each ontology, one for each span detection method using their default parameters. Ten-fold cross-validation was employed for the CRF method but only two-fold cross-validation for the deep learning method, as the training for deep learning takes a long time, and no meaningful differences in performance were observed (see Results). Code, parameter settings and other materials necessary to reproduce these findings are available at https://github.com/UCDenver-ccp/emnlp2017-bilstm-cnn-crf.

*Normalization* is approached as a sequence-to-sequence mapping task, mapping characters in the text spans identified in the previous step to the sequence of characters that constitute an ontology class identifier (e.g., GO:0032502). The deep learning architecture that generated the best results was the OpenNMT system, which implements stacked Bi-LSTM with attention models, and learns condensed vector representations of characters from the training data. The model architecture, as depicted in Figure 1, is character-based, which means that both the encoder and the decoder process one character at a time.

**Figure 1:**
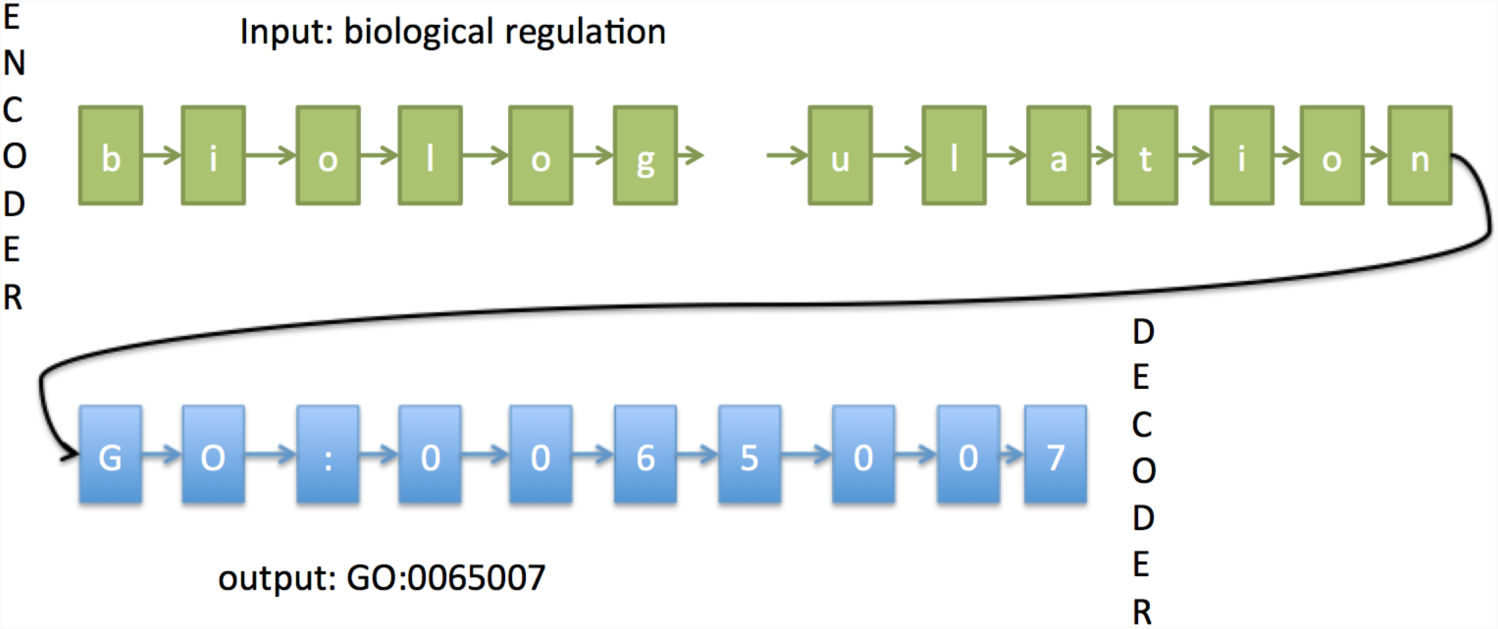
High level view of the character-based encoder-decoder model for concept normalization.

Figure 2 details one layer of the sequence-to-sequence LSTM model. It has four main components: an encoder, a decoder, an attention mechanism and a softmax layer, which are depicted in green, orange, blue and gray boxes, respectively. The OpenNMT implementation stacks multiple layers of encoders, attention mechanims and decoders before the softmax layer at the top.

The size of the input sequence for the encoder is the length of the longest span in the training data, which could be up to 100 characters depending on the ontology. Any input that is shorter is padded with null characters at the end. The size of the output sequence is set to be the number of characters in the ontology identifier, which ranges from 10-17 characters depending on the ontology. CRAFT extension classes were assigned identifiers that start with the ontology names plus “_EXT_” followed by integers assigned sequentially.

To understand the contributions of the various aspects of this architecture to overall performance, the one layer encoder-decoder architecture, which is shown in Figure 2 that takes one-hot-encoding representation of the textual mentions, is also trained.

**Figure 2:**
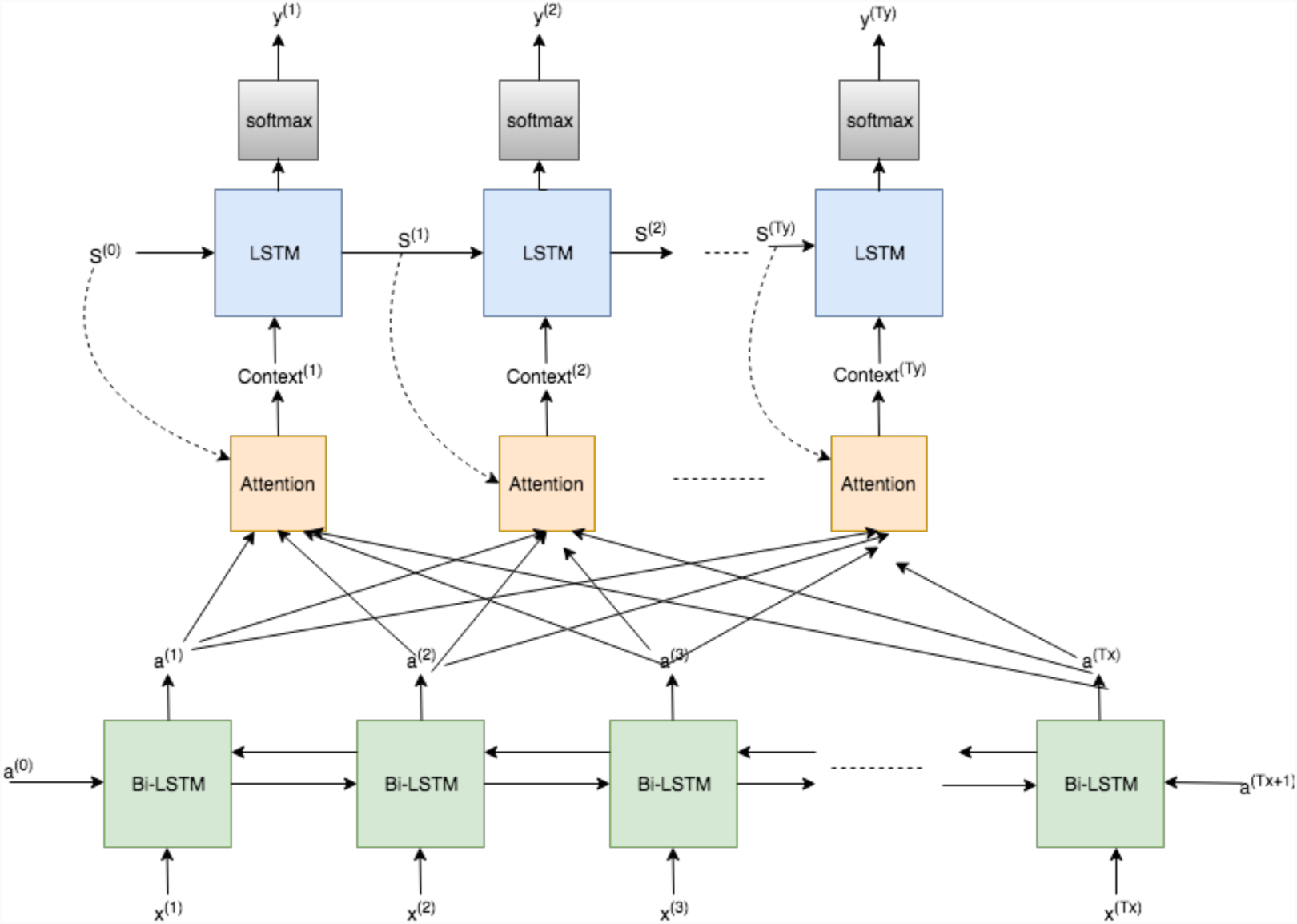
sequence-to-sequence LSTM model with attention.

One of the OBO Foundry identifier policy is that identifiers should not have semantic content (http://www.obofoundry.org/id-policy.html). To explore the potential role of semantics in the outputs, we also mapped the numeric portion the ontology class identifiers to randomly generated digit strings of the same length, and trained models using these as the output representation.

CRAFT, the source of all training and testing data, does not include mentions of all the classes in all of the ontologies. (Actually, most of the classes are not mentioned in the corpus, though this is partly due to the fact that many very specific concepts represented by classes with long, complex names appear only rarely in the biomedical literature.) Normalization performance on classes not included in the training data is likely to be worse than for those in the training data. It is not possible to evaluate performance for such terms directly, as there is no gold standard. However, it is straightforward to generate synthetic training data for normalization based on the labels and synonyms for all of the ontology terms not present in CRAFT. We have no gold standard to evaluate these synthetic data, but training is likely to improve performance on these classes not used in any CRAFT annotations to be similar to performance on classes in the training set. However, inclusion of many thousand additional classes could potentially reduce performance on classes used in CRAFT, as they introduce additional ambiguity and increase the density of the output space, so performance on the CRAFT data after training on all classes was also evaluated.

All code and data are freely available under a CC-BY license. The code is available at https://github.com/UCDenver-ccp/concept-mapping-using-neural-machine-translation-with-attention and the data at https://github.com/UCDenver-ccp/CRAFT.

## Results

Evaluation was with CRAFT as the gold standard. Our best-performing system using the exact span match criterion was usually the combination of a CRF for span detection with a 1-layer sequence-to-sequence model, shown as CRF+1-layer seq2seq for CRAFT v.1, and with an OpenNMT sequence-to-sequence model, shown as CRF+OpenNMT for CRAFT v.3 in Table 1. The differences in the performance of the normalization systems (i.e., the 1-layer seq2seq model performing best for CRAFT v.1, and the OpenNMT being the best over all for CRAFT v.3) could be due to the number of the annotations. There are a larger number of annotations in CRAFT v.3 than CRAFT v.1, and OpenNMT being a larger and deeper network might be doing well with larger training data.

**Table 1:**
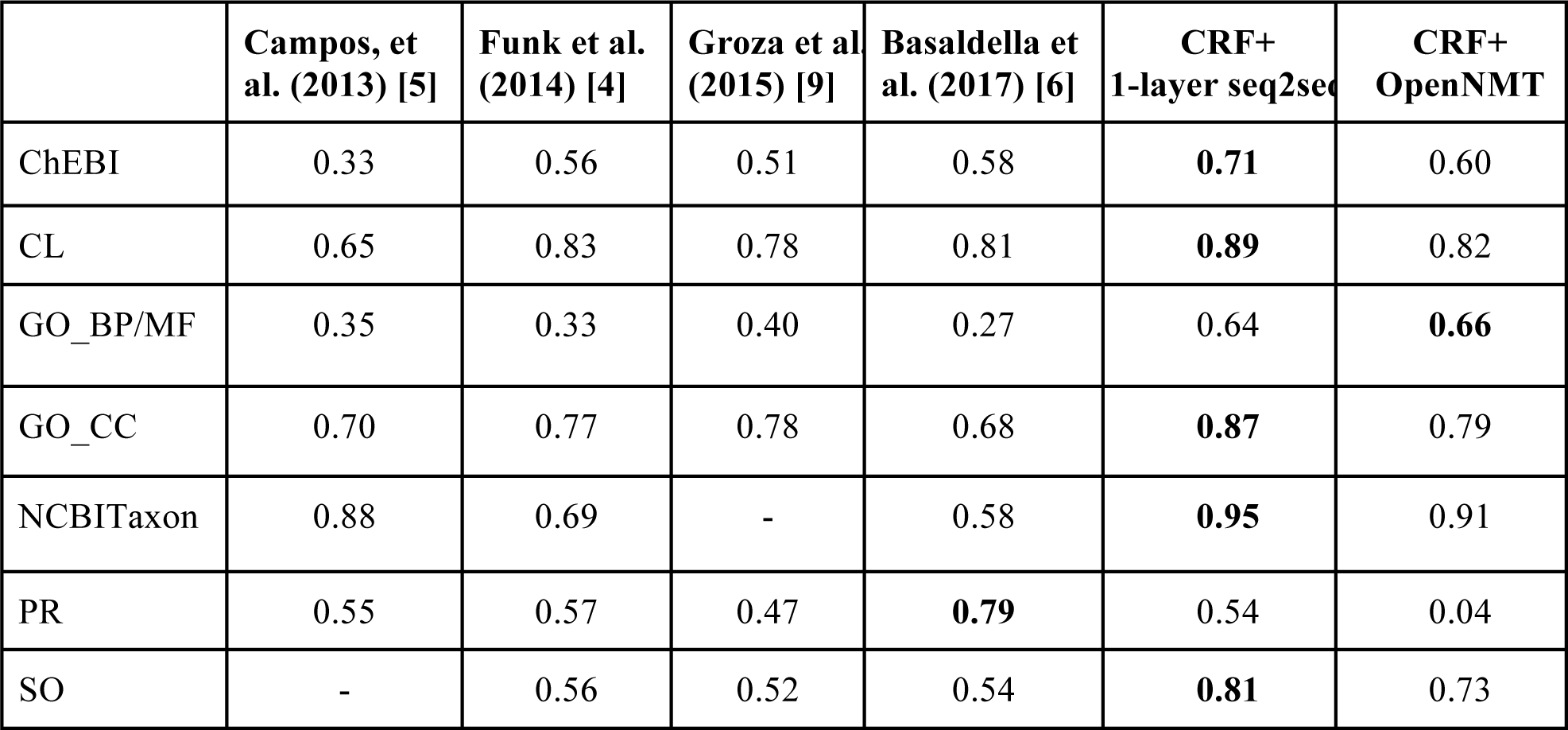
Comparison of end-to-end F1-scores on CRAFT v.1 using an exact span match criterion. The highest result for each CRAFT concept annotation set is in bold font. A hyphen (-) indicates that the given study did not publish results for the given CRAFT concept annotation set.

|  | Campos, et al. (2013) [5] | Funk et al. (2014) [4] | Groza et al (2015) [9] | Basaldella et al. (2017) [6] | CRF+ 1-layer seq2seq | CRF+ OpenNMT |
| --- | --- | --- | --- | --- | --- | --- |
| ChEBI | 0.33 | 0.56 | 0.51 | 0.58 | **0.71** | 0.60 |
| CL | 0.65 | 0.83 | 0.78 | 0.81 | **0.89** | 0.82 |
| GO_BP/MF | 0.35 | 0.33 | 0.40 | 0.27 | 0.64 | **0.66** |
| GO_CC | 0.70 | 0.77 | 0.78 | 0.68 | **0.87** | 0.79 |
| NCBITaxon | 0.88 | 0.69 | - | 0.58 | **0.95** | 0.91 |
| PR | 0.55 | 0.57 | 0.47 | **0.79** | 0.54 | 0.04 |
| SO | - | 0.56 | 0.52 | 0.54 | **0.81** | 0.73 |

### End-to-end system evaluation

The primary evaluation is end-to-end concept recognition performance, that is, the combination of span detection followed by normalization. Table 1 compares the CRF+1-layer seq2seq vs. CRF+OpenNMT systems against four recently published high performance concept recognition systems, requiring exact match for spans. Evaluations were performed on a holdout test set of 7 randomly selected articles (∼10%) of the CRAFT v.1 corpus. The CRF+1-layer seq2seq system outperforms the existing state-of-the-art concept normalization systems for all ontologies except the Protein Ontology (PR) often reducing error rates by 50% or more. Basaldella et al. remains the state of the art for Protein Ontology classes. The CRF+OpenNMT system also outperforms the other systems for most ontologies and is the best-performing system for the GO_BP/MF annotation set.

Performance varies slightly when using the deep learning span detection approach (See Table 4 for details), but in all cases the differences were less than small and unlikely to be meaningful in general. Since the CRF span detection method is much faster to train than the already widely used deep learning approach and performs nearly identically, it is used in all the end-to-end comparisons.

As CRAFT v.3 has been released only recently, there are no published performance statistics for its updated concept annotation sets, nor are there any published results for concept recognition for classes from the Molecular Process Ontology (MOP) or the Uberon anatomical ontology or of CRAFT extension classes (all of which are new in version 3.0). Although there was reason to believe the extension classes might have been more difficult (see discussion above), Table 2 shows the F1 score of the CRF+OpenNMT system remains outstanding. For tasks requiring only the correct identification concepts, the exact span match criterion may be too strict, so Table 3 shows performance for a criterion requiring only overlapping spans (of at least one character) along with the correct class, which is likely to be the best indication of performance in biomedical text generally. In all cases, allowing overlap rather than exact matching caused either a slight increase or no change in performance. Intriguingly, allowing inexact matches generally increased both precision and recall, although in all cases the increases were very small.

**Table 2:** Comparison of end-to-end F1-scores on CRAFT v.3 using an exact span match criterion. The highest result for each CRAFT concept annotation set is in bold font. A hyphen (-) indicates that the given study did not publish results for the given CRAFT concept annotation set.

|  | CRAFT v.3 proper OBO classes |  | CRAFT v.3 proper OBO classes plus extension classes |  |
| --- | --- | --- | --- | --- |
|  | CRF+ 1-layer seq2seq | CRF+ OpenNMT | CRF+ 1-layer seq2seq | CRF+ OpenNMT |
| ChEBI | 0.63 | 0.66 | 0.68 | 0.71 |
| CL | 0.74 | 0.76 | 0.80 | 0.80 |
| GO_BP | 0.74 | 0.76 | 0.66 | 0.80 |
| GO_CC | 0.73 | 0.72 | 0.87 | 0.87 |
| GO_MF | - | - | 0.69 | 0.69 |
| MOP | - | - | 0.75 | 0.75 |
| NCBITaxon | 0.95 | 0.95 | 0.96 | 0.96 |
| PR | 0.07 | 0.23 | 0.36 | 0.47 |
| SO | 0.89 | 0.90 | 0.78 | 0.80 |
| UBERON | 0.77 | 0.79 | 0.72 | 0.79 |

**Table 3:** Performance of CRF+OpenNMT concept recognition on CRAFT version 3 ontologies, including extensions. Overlapping vs. Exact refers to span match.

|  | Overlapping span match |  |  | Exact span match |  |  |
| --- | --- | --- | --- | --- | --- | --- |
|  | P | R | F1 score | P | R | F1 score |
| ChEBI | 0.88 | 0.61 | 0.72 | 0.87 | 0.60 | 0.71 |
| CL | 0.86 | 0.77 | 0.81 | 0.85 | 0.76 | 0.80 |
| GO_BP | 0.86 | 0.77 | 0.81 | 0.85 | 0.75 | 0.80 |
| GO_CC | 0.94 | 0.84 | 0.88 | 0.92 | 0.82 | 0.87 |
| GO_MF | 0.86 | 0.59 | 0.70 | 0.85 | 0.58 | 0.69 |
| MOP | 0.95 | 0.61 | 0.75 | 0.95 | 0.61 | 0.75 |
| NCBITaxon | 1.0 | 0.93 | 0.96 | 0.99 | 0.92 | 0.96 |
| PR | 0.65 | 0.38 | 0.48 | 0.64 | 0.38 | 0.47 |
| SO | 0.81 | 0.76 | 0.79 | 0.83 | 0.77 | 0.80 |
| UBERON | 0.81 | 0.76 | 0.79 | 0.81 | 0.76 | 0.79 |

### Performance Analysis

As noted above, span detection method made relatively little difference in end-to-end performance. Table 4 compares the end-to-end performance with a CRF versus deep learning for span detection (using the overlapping criterion) on the CRAFT v3.0 corpus. The differences in performance are small and vary in direction, showing no particular trend toward either method.

The OBO Foundry ID principles^2^ discourage the use of identifiers that have semantic content. However, the performance of the OpenNMT concept normalization system suggests that the network might be finding semantic information in those identifiers. To test this hypothesis, the system was retrained with randomly generated identifiers for the outputs.

As shown in Table 5, performance on randomized identifiers dropped modestly for several ontologies, indicating that there is indeed semantic information encoded in IDs. This evaluation is on gold textual span using the accuracy metrics for multi-label classification^3^; however, the difference is not clearly seen when the model is evaluate in automatically extracted textual span as shown in Table 6. We computed Table 5 using micro averaged F1 scores but results remained the same. Discussion with a Gene Ontology Consortium director (personal communication) suggests a possible mechanism: Specialist groups are convened from time to time to expand or revise particular topics in an ontology; these groups are given a set of consecutive identifiers to use as they see fit. As a result, consecutive identifiers have a greater than random chance of being semantically related, and the OpenNMT normalization network appears to have found this signal.

**Table 4:**
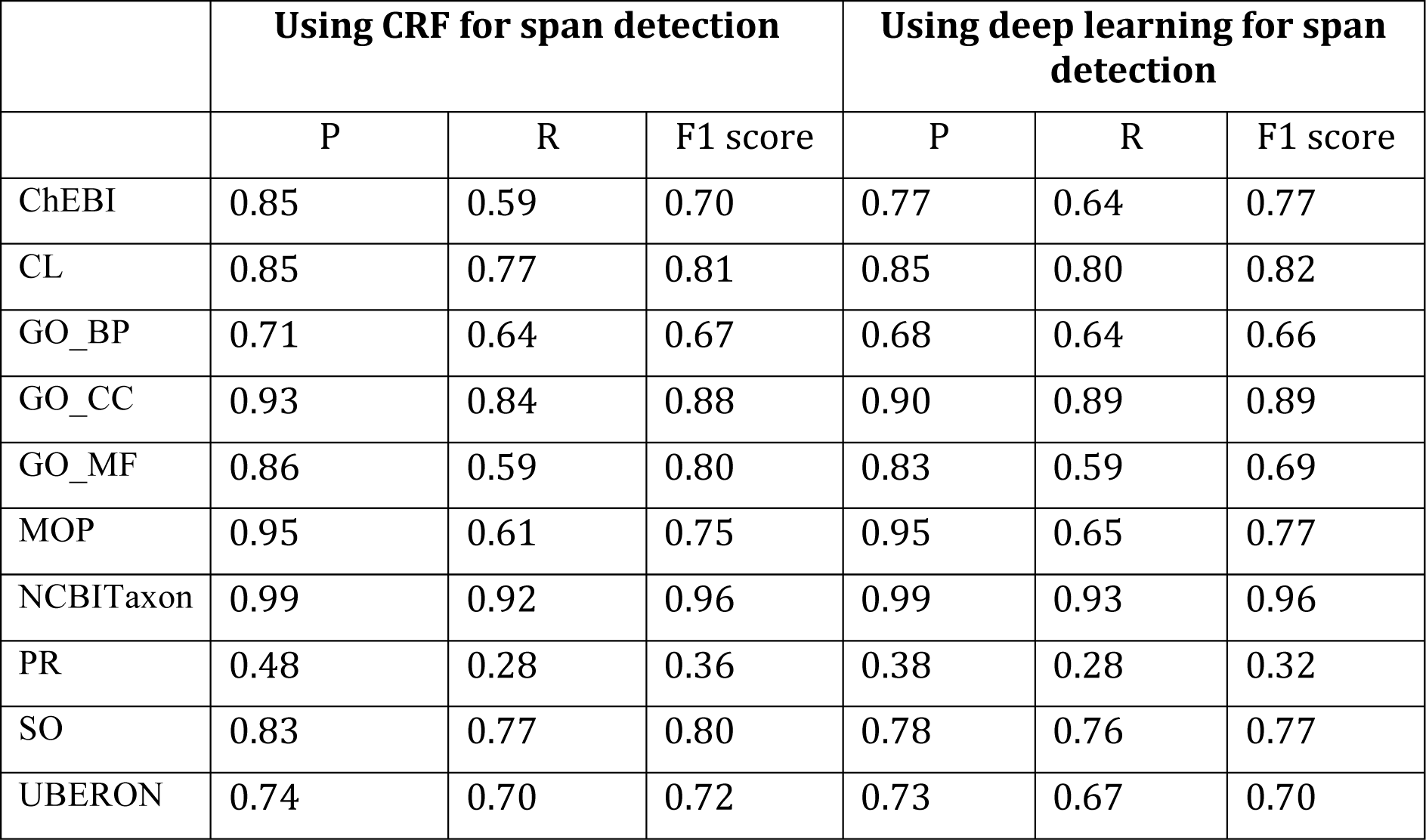
End-to-end performance is modestly affected by span detection method. Evaluation is against CRAFT v.3.

|  | Using CRF for span detection |  |  | Using deep learning for span detection |  |  |
| --- | --- | --- | --- | --- | --- | --- |
|  | P | R | F1 score | P | R | F1 score |
| ChEBI | 0.85 | 0.59 | 0.70 | 0.77 | 0.64 | 0.77 |
| CL | 0.85 | 0.77 | 0.81 | 0.85 | 0.80 | 0.82 |
| GO_BP | 0.71 | 0.64 | 0.67 | 0.68 | 0.64 | 0.66 |
| GO_CC | 0.93 | 0.84 | 0.88 | 0.90 | 0.89 | 0.89 |
| GO_MF | 0.86 | 0.59 | 0.80 | 0.83 | 0.59 | 0.69 |
| MOP | 0.95 | 0.61 | 0.75 | 0.95 | 0.65 | 0.77 |
| NCBITaxon | 0.99 | 0.92 | 0.96 | 0.99 | 0.93 | 0.96 |
| PR | 0.48 | 0.28 | 0.36 | 0.38 | 0.28 | 0.32 |
| SO | 0.83 | 0.77 | 0.80 | 0.78 | 0.76 | 0.77 |
| UBERON | 0.74 | 0.70 | 0.72 | 0.73 | 0.67 | 0.70 |

**Table 5:** Comparison of performance between OBO assigned identifiers and randomly assigned identifiers on CRAFT v.3 proper obo classes on gold textual mentions.

|  | With OBO assigned IDs in accuracy | With randomly assigned IDs in accuracy |
| --- | --- | --- |
| ChEBI | 74.58 | 71.56 |
| CL | 82.44 | 82.68 |
| GO_BP | 86.82 | 87.38 |
| GO_CC | 89.96 | 86.62 |
| NCBITaxon | 95.79 | 96.07 |

<sup>2</sup> http://www.obofoundry.org/principles/fp-003-uris.html

|  |  |  |
| --- | --- | --- |
| PR | 42.81 | 42.58 |
| SO | 96.09 | 96.34 |
| UBERON | 85.33 | 84.09 |

**Table 6:** Comparison of performance between OBO assigned identifiers and random identifiers on automatically extracted textual spans. Evaluation is on CRAFT v.3 proper OBO classes.

|  | With OBO assigned IDs |  |  |  |  |  | With randomly assigned IDs |  |  |  |  |  |
| --- | --- | --- | --- | --- | --- | --- | --- | --- | --- | --- | --- | --- |
|  | Overlapping span match |  |  | Exact span match |  |  | Overlapping span match |  |  | Exact span match |  |  |
|  | P | R | F1 score | P | R | F1 score | P | R | F1 score | P | R | F1 score |
| ChEBI | 0.93 | 0.51 | 0.66 | 0.93 | 0.51 | 0.66 | 0.92 | 0.51 | 0.65 | 0.92 | 0.51 | 0.65 |
| CL | 0.84 | 0.71 | 0.77 | 0.83 | 0.70 | 0.76 | 0.82 | 0.70 | 0.75 | 0.82 | 0.69 | 0.75 |
| GO_BP | 0.85 | 0.70 | 0.76 | 0.85 | 0.70 | 0.76 | 0.84 | 0.69 | 0.76 | 0.83 | 0.69 | 0.75 |
| GO_CC | 0.87 | 0.65 | 0.74 | 0.84 | 0.63 | 0.72 | 0.87 | 0.65 | 0.75 | 0.85 | 0.64 | 0.73 |
| GO_MF | - | - | - | - | - | - | - | - | - | - | - | - |
| MOP | - | - | - | - | - | - | - | - | - | - | - | - |
| NCBITaxon | 1 | 0.92 | 0.96 | 0.99 | 0.92 | 0.95 | 1 | 0.92 | 0.96 | 0.99 | 0.92 | 0.95 |
| PR | 0.44 | 0.15 | 0.23 | 0.44 | 0.15 | 0.23 | 0.45 | 0.15 | 0.22 | 0.45 | 0.15 | 0.22 |
| SO | 0.93 | 0.88 | 0.90 | 0.93 | 0.88 | 0.90 | 0.93 | 0.88 | 0.90 | 0.93 | 0.88 | 0.90 |
| UBERON | 0.83 | 0.56 | 0.79 | 0.83 | 0.76 | 0.79 | 0.83 | 0.76 | 0.80 | 0.83 | 0.76 | 0.79 |

The OpenNMT system provides more functionality than a simple LSTM architecture, adding vector encodings for characters, and a stacked (multilayer) LSTM architecture. To evaluate how much these features contribute to performance, the 1-layer sequence-to-sequence architecture depicted in Figure 2 with a one-hot encoding for characters was trained on the same data, with results shown in Table 7. In no case did the additional features in the OpenNMT system cause performance degradation, and in some cases (especially GO_BP) they provided significant increases in performance.

**Table 7:** The stacked architecture and character vector encodings in OpenNMT improve performance over a 1-layer sequence to sequence architecture with one-hot encoding against CRAFT v.3 proper OBO class plus extension classes

|  | CRF +OpenNMT |  |  |  |  |  | CRF+1-layer seq2seq |  |  |  |  |  |
| --- | --- | --- | --- | --- | --- | --- | --- | --- | --- | --- | --- | --- |
|  | Overlapping span match |  |  | Exact span match |  |  | Overlapping span match |  |  | Exact span match |  |  |
|  | P | R | F1 score | P | R | F1 score | P | R | F1 score | P | R | F1 score |
| ChEBI | 0.88 | 0.61 | 0.72 | 0.87 | 0.6 | 0.71 | 0.85 | 0.59 | 0.7 | 0.82 | 0.57 | 0.68 |
| CL | 0.86 | 0.77 | 0.81 | 0.85 | 0.76 | 0.8 | 0.85 | 0.77 | 0.81 | 0.84 | 0.76 | 0.8 |
| GO_BP | 0.86 | 0.77 | 0.81 | 0.85 | 0.75 | 0.8 | 0.71 | 0.64 | 0.67 | 0.7 | 0.63 | 0.66 |
| GO_CC | 0.94 | 0.84 | 0.88 | 0.92 | 0.82 | 0.87 | 0.93 | 0.84 | 0.88 | 0.92 | 0.83 | 0.87 |
| GO_MF | 0.86 | 0.59 | 0.7 | 0.85 | 0.58 | 0.69 | 0.86 | 0.59 | 0.7 | 0.86 | 0.58 | 0.69 |
| MOP | 0.95 | 0.61 | 0.75 | 0.95 | 0.61 | 0.75 | 0.95 | 0.61 | 0.75 | 0.95 | 0.61 | 0.75 |
| NCBITaxon | 1 | 0.93 | 0.96 | 0.99 | 0.92 | 0.96 | 0.99 | 0.92 | 0.96 | 0.99 | 0.92 | 0.96 |
| PR | 0.65 | 0.38 | 0.48 | 0.64 | 0.38 | 0.47 | 0.48 | 0.28 | 0.36 | 0.48 | 0.28 | 0.36 |
| SO | 0.81 | 0.76 | 0.79 | 0.83 | 0.77 | 0.8 | 0.83 | 0.77 | 0.8 | 0.81 | 0.75 | 0.78 |
| UBERON | 0.81 | 0.76 | 0.79 | 0.81 | 0.76 | 0.79 | 0.74 | 0.7 | 0.72 | 0.74 | 0.69 | 0.72 |

One issue with machine-learning-based approaches to concept recognition in text is that such systems are unlikely to recognize classes that are not present in the training data. CRAFT is the most extensive manually annotated corpus for most of these ontologies, but there are many classes in the ontologies that do not appear in CRAFT at all; for example, the NCBI Taxonomy contains more than 400,000 taxa, fewer than 100 of which appear in CRAFT. However, as previously stated, this phenomenon is also partly due to the fact that many specific concepts represented by ontology classes with long, complex names appear only rarely in the biomedical literature. The separation of span detection from concept normalization allows a straightforward approach to addressing this problem: we can expand the training set for the OpenNMT system with synthetic training data of all of the class names and synonyms from all of the ontologies, with spans defined by the length of the name or synonym. While the lack of gold-standard data means it is impossible to evaluate the performance of such an approach, Table 8 shows how training on all class names and synonyms affects performance on the terms that do appear in CRAFT. Unsurprisingly, performance decreases when training with many thousands of additional classes. The changes are mostly modest, except for ChEBI and PR where the large number of new terms caused a drop in performance from F1 of 0.66 to 0.54 for ChEBI and from 0.23 to 0.06 for PR. The NCBI Taxonomy has a huge number of classes in the synthetic data, but performance with that synthetic data remains good with an F1 of 0.87. The performance on CRAFT terms with the synthetic data is probably the most reasonable estimates of the ability of the approach to detect any ontology term in biomedical text. Even the reduced performance on the CRAFT classes with the additional training data is significantly better than the current state of the art (except in PR). It would also be possible to use synonym expansion techniques [41] to generate additional training examples.

**Table 8:** End-to-end performance on CRAFT v.3 proper OBO classes with synthetic data reflecting all ontology terms and synonyms for each ontology. GO_MF and MOP omitted due to low numbers of annotations in CRAFT.

|  | CRAFT v.3 without synthetic data |  |  |  |  |  | CRAFT v.3 + synthetic data |  |  |  |  |  |
| --- | --- | --- | --- | --- | --- | --- | --- | --- | --- | --- | --- | --- |
|  | Overlapping span match |  |  | Exact span match |  |  | Overlapping span match |  |  | Exact span match |  |  |
|  | P | R | F1 score | P | R | F1 score | P | R | F1 score | P | R | F1 score |
| ChEBI | 0.93 | 0.51 | 0.66 | 0.93 | 0.51 | 0.66 | 0.76 | 0.42 | 0.54 | 0.76 | 0.42 | 0.54 |
| CL | 0.84 | 0.71 | 0.77 | 0.83 | 0.7 | 0.76 | 0.83 | 0.7 | 0.76 | 0.82 | 0.7 | 0.75 |
| GO_BP | 0.85 | 0.7 | 0.77 | 0.85 | 0.7 | 0.76 | 0.82 | 0.68 | 0.74 | 0.8 | 0.65 | 0.72 |
| GO_CC | 0.87 | 0.65 | 0.74 | 0.84 | 0.63 | 0.72 | 0.85 | 0.65 | 0.74 | 0.81 | 0.61 | 0.7 |
| GO_MF | - | - | - | - | - | - | - | - | - | - | - | - |
| MOP | - | - | - | - | - | - | - | - | - | - | - | - |
| NCBITaxon | 1 | 0.92 | 0.96 | 0.99 | 0.92 | 0.95 | 0.91 | 0.84 | 0.87 | 0.91 | 0.84 | 0.87 |
| PR | 0.44 | 0.15 | 0.23 | 0.44 | 0.15 | 0.23 | 0.12 | 0.04 | 0.06 | 0.12 | 0.04 | 0.06 |
| SO | 0.93 | 0.88 | 0.9 | 0.93 | 0.88 | 0.9 | 0.93 | 0.88 | 0.9 | 0.93 | 0.88 | 0.9 |
| UBERON | 0.83 | 0.76 | 0.79 | 0.83 | 0.76 | 0.79 | 0.83 | 0.76 | 0.8 | 0.83 | 0.76 | 0.79 |

### Discussion and Conclusions

Modeling the concept recognition task as a sequence-to-sequence machine translation problem enable the application of modern deep learning technology to the task, resulting in a striking improvement in performance over previous methods. Recognition in full text biomedical journal articles of biomedical ontology classes from nearly any ontology can now be achieved with an F1 score of 0.7 or better, opening up a variety of new applications in information retrieval and information extraction in biomedical text.

The improvements in performance were almost entirely due to improvements in concept normalization. Deep learning methods for span detection were equivalent in performance to traditional conditional random field methods. The conception of the concept normalization problem as sequence-to-sequence mapping and the Bi-LSTM architecture are responsible for most of the performance, but adding additional LSTM layers and a vector encoding for characters are also significant. There is some evidence that the deep learner is capturing semantic information embedded in the ontology class IDs as well.

As natural training data is limited to the concepts used in CRAFT annotations, the addition of synthetic training data, class names and synonyms, to the normalization step has the potential to improve recall on class not in CRAFT. Evaluation on the performance of OpenNMT with added synthetic training data on CRAFT shows a decrease in performance for ChEBI class, but nearly unchanged performance for the other ontologies.

## Appendix A: list of acronyms

|  |  |
| --- | --- |
| CRFs | Conditional Random Fields |
| RNN | Recurrent Neural Network |
| LSTM | Long Short Term Memory |
| Bi-LSTM | Bidirectional Long Short Term Memory |

## Footnotes

1 http://www.biocreative.org/

2 http://www.obofoundry.org/principles/fp-003-uris.html

3 https://scikit-learn.org/stable/modules/generated/sklearn.metrics.accuracy_score.html

